# mc-ASTRA maps population-level variation to interpretable multicellular tissue organization processes

**DOI:** 10.64898/2026.09.18.752722

**Authors:** Ricardo O. Ramirez Flores, Klaus Sebastian Augusto Kruger Serrano, Francisca Gaspar Vieira, Loan Vulliard, Julio Saez-Rodriguez

## Abstract

Single-cell and spatial technologies increasingly enable detailed molecular profiling of tissues across biological contexts, creating opportunities to relate variation across samples to the multicellular processes that give rise to differences in tissue organisation and function. However, existing tools typically treat these aspects separately, either representing variation across samples or interpreting specific molecular and spatial features. To bridge this gap, we developed multicellular Analysis of Sample Tissue Representations and Associations (mc-ASTRA; https://github.com/saezlab/mc-astra), an open-source Python package for constructing maps of tissue-state variability that combine diverse tissue descriptors, including molecular, compositional and spatial features, together with complementary sample-level information such as clinical measurements or technical covariates, and biological prior knowledge of processes of interest. By combining available flexible semi-supervised factor models with established tools for the inference of biological processes from omics data, mc-ASTRA makes these tissue-state maps interpretable by quantifying how different tissue descriptors contribute to variation across samples and by resolving the specific coordinated multicellular programs underlying these differences. We illustrate the capabilities of mc-ASTRA across multiple single-cell and spatial omics datasets to infer multicellular programs that are contextualized by technical and biological prior knowledge and to describe distinct trajectories of tissue remodeling. mc-ASTRA provides a framework for building interpretable sample maps that connect tissue organization and multicellular mechanisms with variation observed across samples.

**Graphical Abstract:** 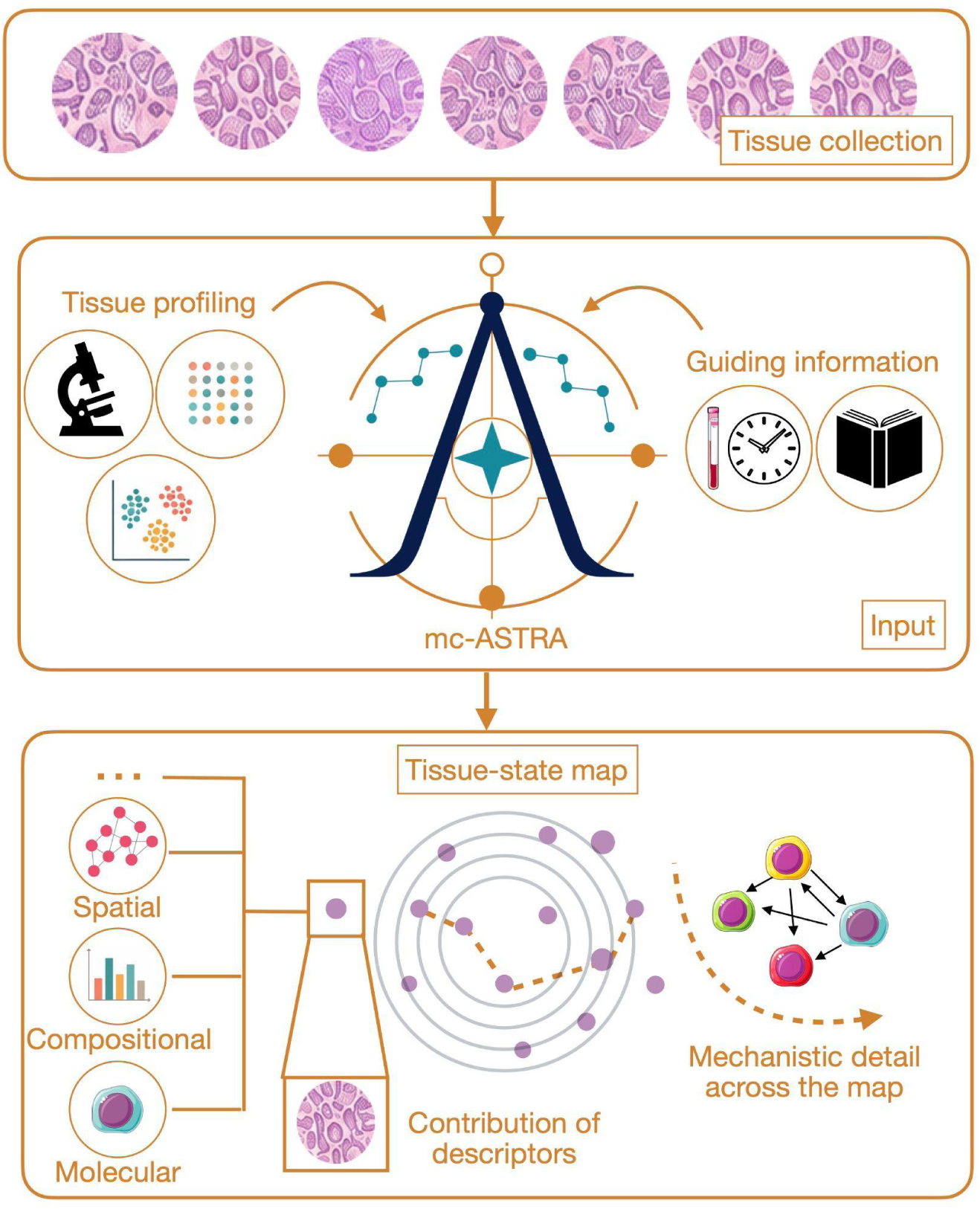

## 1. Introduction

Modern biological and medical studies increasingly rely on single-cell and spatial technologies to characterize tissues at high resolution, revealing variation in molecular activity, cell type compositions and spatial organization^1^. This abundance of profiling creates an opportunity to explore tissue variability beyond the one reflected in conventional annotations (e.g. patient diagnoses or experimental conditions such as diet). For example, samples sharing the same annotation can differ substantially in their underlying tissue biology ^2–4^, while samples assigned to different groups can exhibit similar cellular and molecular states^2,5,6^. To better represent this continuous variation in tissue biology, each sample can be positioned in a latent space by combining multiple descriptors of its tissue state. These latent spaces, here referred to as tissue-state maps, are intended to retain biological and technical information, accommodate new data, and remain interpretable by grounding sample representations in specific tissue components and multicellular biological processes that can support mechanistic and actionable models.

Methodologically speaking, tissue-state maps built from molecular data at distinct resolutions are an example of multimodal representation learning tasks that integrate and interpret different sources of information simultaneously. As such, they face two general challenges: what information to integrate, and how to construct the latent space? In the context of tissue-state maps built from single-cell molecular data, tissue samples can be represented through complementary tissue descriptors capturing different aspects of multicellular organisation, including, among others, molecular features (e.g., gene expression), composition (e.g., cell-state abundance), spatial organisation (e.g., cell-state colocalisation or neighbourhood relationships), and functional activity (e.g., cell-cell communication or pathway activation)^7^. Statistical methods used to reconstruct tissue-state maps should be able to handle these distinct tissue descriptors, either incrementally or simultaneously, and to associate each of them with disease manifestation.

Current methods to reconstruct tissue-state maps use two broad strategies (Supplemental Table 1). Some make their primary objective the definition of pairwise relationships between samples, typically through similarity or distance metrics, and later use these relationships to construct a low-dimensional embedding ^8–12^. Others formulate tissue-state mapping as a reconstruction problem of the original multimodal data and represent either single-cell measurements or patient-level transformed data in fewer dimensions using probabilistic generative models ^13–15^, foundation models ^16,17^, or matrix or tensor factorization ^18–23^. Most current methods, however, fix a single representation of tissue descriptors, such as gene expression or cellular composition ^24^, and do not interpret the latent space in terms of multicellular function. Recent evaluation frameworks have facilitated direct comparison between different strategies for building tissue-state maps^24,25^, but their reliance on reported sample labels assumes that these indicators explain a substantial proportion of the variance in tissue descriptors and that these annotations are done correctly. In most cases, whether this assumption holds is unknown and is one reason why molecular profiling is performed in the first place. For example, the chronological age of a patient does not necessarily reflect the biological age for every cell-type^26^ and clinical and pathological diagnostic adjudication processes can lead to different classifications of disease etiology^27^. Moreover, because available methods combine different computational strategies with different tissue descriptors, comparisons based only on predictive performance are difficult to interpret since improved performance may arise from the model, the chosen representation, or both.

We previously proposed repurposing probabilistic group factor models to build tissue-state maps from single-cell and spatial data using multicellular factor analysis, and showed that their flexibility in data handling and model specification allowed multicellular gene-expression programs to be studied together with other tissue or patient descriptors across multiple cohorts^19^. Several studies have shown the usefulness of multicellular factor analysis for multicellular disease description^28^, meta-analysis^6,29^, and association with clinical outcomes in large patient cohorts across omics technologies^30–32^. However, a limitation of the current unsupervised model was the lack of flexibility for adding structure to the learning of the tissue-state maps using prior knowledge, either from complementary sample annotations (e.g. clinical or demographic data of a patient cohort) or from biological processes of interest (e.g. perturbation gene expression signatures). Recent methodological and computational extensions of group factor models have substantially expanded the range of model structures that can be specified, including the incorporation of structured priors over samples and features^33–36^. However, applying these models to tissue-state analysis requires an additional framework that transforms single-cell and spatial measurements into interpretable sample-level representations and connects the resulting latent factors to downstream analyses to infer multicellular processes.

To fill this gap we present multicellular Analysis of Sample Tissue Representations and Associations (mc-ASTRA; https://github.com/saezlab/mc-astra), an open-source Python package that provides the framework to define, construct and interpret multicellular tissue-state maps end-to-end. mc-ASTRA facilitates the integration of cellular composition, spatial organization, and gene expression from emerging single-cell and spatial datasets, while allowing flexible models to generate biologically relevant interpretable tissue-state representations and assess the contributions of distinct tissue descriptors. Across diverse single-cell and spatial applications, mc-ASTRA captured known and latent sources of tissue variation, contextualized molecular prior knowledge from perturbation data, resolved continuous disease-associated tissue states, and identified spatiotemporal multicellular programs linking structural and molecular changes during damage and recovery.

## 2. Results

### 2.1 mc-ASTRA’s architecture

mc-ASTRA’s primary objective is to derive potential multicellular processes that describe the variability of a given sample collection (e.g. patient cohort) profiled molecularly at the single-cell or spatial level with optional additional descriptors (e.g. clinical data, pathology features, etc.) by combining tissue descriptor generators, multicellular factor analysis, and downstream functional analyses (Figure 1). To perform multicellular and multimodal integration tasks, mc-ASTRA relies on the flexibility of probabilistic group factor analysis implemented in MOFA-FLEX ^33^ to build a latent space, here referred to as tissue-state map, from a multi-view representation of a sample collection. In this representation, the same samples are described by multiple sets of features, or views, each capturing a distinct tissue descriptor, such as cell-type composition or the gene-expression profile of a specific cell population. Within this map, each sample is positioned according to its combination of tissue characteristics, while distances between samples capture similarities and differences that can support stratification, ordering along latent dimensions to represent tissue remodeling processes, and trajectory inference. Given the linear nature of factor models, the tissue-state map can be interpreted in terms of the original variables and be used to project new sample collections. In addition, the model’s reconstruction errors can help identify relevant axes of variation and the different tissue descriptors contributing to these differences. In combination with functional transcriptomics tools, latent variables can be transformed into potential mechanistic hypotheses of multicellular regulation, for example, by building cell to cell networks capturing coordination and ligand-receptor usage (Figure 1).

**Figure 1.**
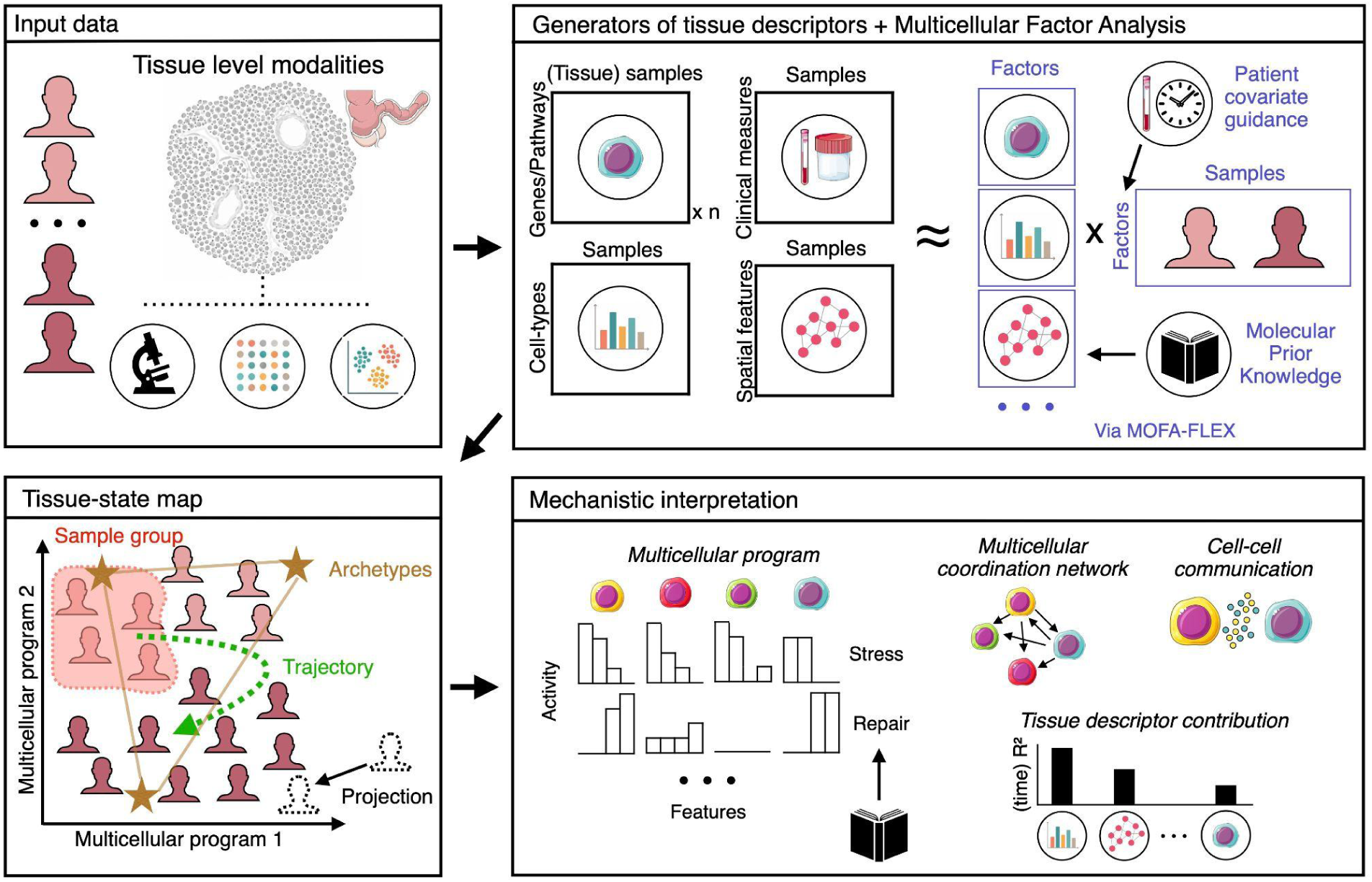
Multicellular Analysis of Sample Tissue Representations and Associations (mc-ASTRA) objectives and functionalities. mc-ASTRA provides an end-to-end framework for the construction and multicellular interpretation of tissue-state maps from multimodal data. Focused primarily on the integration of single-cell, spatial, and clinical data, mc-ASTRA combines multiple tools for the definition of tissue descriptors, and tissue-state map construction and interpretation. This integration of functionalities allows for the construction of more flexible tissue-state maps that take into account distinct descriptors (e.g., spatial cellular organization) and can be guided by additional prior knowledge, such as sample covariates (e.g., time or clinical measures) or feature annotations (e.g., pathway activity gene sets). Tissue-state map representations are stored as *anndata*^41^, a commonly used data structure for single-cell analysis, allowing users to apply multiple methodologies for clustering, trajectory inference, projection, or archetypal analysis already available, for example, through the *scverse^42^* ecosystem. mc-ASTRA provides multiple functionalities that facilitate the interpretation of tissue-state maps as multicellular programs, coordinated activities across cell types that can be associated with spatial or compositional tissue-remodelling processes. Cell-type specific and multicellular active molecular processes can be inferred from these programs, as well as potential coordination and communication processes. Moreover, the model’s reconstruction errors can be used to quantify the importance of specific tissue descriptors in explaining the variance associated to specific sample covariates. Colon and cell images provided by Servier Medical Art (https://smart.servier.com), licensed under CC BY 4.0.

The basic workflow assumes that the tissue descriptors of a sample collection can be transformed into a multi-view sample-level representation. In the case of single-cell data, for example, these views are usually the pseudobulk expression profile of each cell type defined by the researchers. However, these representations are not limited to pseudobulk representations, since they could be compositional views, spatial summary metrics, pathway activities, or any feature space where there is a quantification per patient^19,20,32,37,38^. In mc-ASTRA, we take advantage of different established packages for single-cell data analysis to provide unified quality control and analysis utilities to derive multicellular expression, compositional, and spatial views (see Methods).

From the modeling aspect, mc-ASTRA extends multicellular factor analysis^19^ by taking advantage of the flexible modeling extensions implemented in MOFA-FLEX^33^. In its original formulation, multicellular factor analysis relied on group factor models that accommodated multi-view data with different distributions, missing values, and group-aware modeling^39,40^. MOFA-FLEX further extends these models by allowing them to incorporate sample-level covariates^34–36^, such as time and clinical measurements, or by molecular prior knowledge, including signaling pathway information or perturbation signatures. The semi-supervision aspect of the extended factor models facilitates combining targeted with exploratory analyses, which mc-ASTRA uses to address multiple biological questions of multicellular coordination (Results section 2.2). Some specific examples include the quantification of the independence of molecular and structural tissue remodeling processes, the identification of cell-type specific signaling pathway responses to perturbations, the definition of dynamic tissue remodeling processes from time-course data or the association between circulating biomarkers and cell-type specific damage responses in tissues. In addition, the same type of structured regularization can be used to control technical aspects of the experimental design, to integrate distinct sample cohorts, or to contextualize prior-knowledge to the tissue of interest.

mc-ASTRA transforms models built with MOFA-FLEX to commonly used data structures for single-cell data analysis, facilitating their exploration and interoperability with other analysis packages that operate on *anndata^41^*, including those in the *scverse* ecosystem^42^. For example, model results can be directly analyzed using clustering approaches commonly applied in single-cell workflows to identify discrete sample groups, or combined with recent archetypal analysis implementations to explore the multicellular functional trade-offs captured by the latent space and simplify the interpretation of tissue-state maps with multiple programs associated with covariates of interest^43^ (Results section 2.2.3).

Beyond constructing tissue-state maps, the different layers of interpretability provided by factor models are used in mc-ASTRA to characterize multicellular processes underlying sample variation. The latent variables estimated from the data (i.e. factors) can be interpreted as multicellular programs, i.e. coordinated activities of gene expression that can be coupled to spatial and compositional organization. Each factor score per sample is built from a weighted linear combination of the original features across views (i.e. loadings, Figure 1), thus the weight of each feature represents its importance in the multicellular program. Similarly, the variance explained by the model per factor and view (i.e. the coefficient of determination, R², calculated from reconstruction errors) can be used as a proxy of the importance of a tissue descriptor in a multicellular program or to estimate the proportion of total variance of a covariate of interest explained by the model (Methods).

To determine whether variation captured by the factors is associated with clinical or demographic covariates, mc-ASTRA tests associations between factors and sample annotations using linear mixed models. To enable compatibility with the evaluation frameworks for sample representations available in *patpy*, we implemented an adapter that transforms any tissue-state map generated with mc-ASTRA into an object compatible with *patpy*’s functionalities*^25^*. This integration combines information retention metrics used in current benchmarks with metrics of explained variance, providing users with additional tools to identify the tissue features most relevant to a biological or clinical question.

Altogether, mc-ASTRA presents an approach to build and interpret tissue-state maps from multimodal and multicellular data that bridges state-of-the-art data processing tools, prior knowledge, and flexible factor modeling frameworks. This combination of tools allows researchers to infer multicellular coordination processes associated with sample variability from complex experimental designs while having control to add structure to the tissue-state map using external guiding information of samples and features. These capabilities provide a unified framework for modeling and interpreting tissue-state maps, distinguishing mc-ASTRA from approaches based primarily on pairwise sample distances, deep latent embeddings, or more constrained tensor and factor representations (Supplemental Table 1).

### 2.2 Structured supervision and downstream interpretation of tissue-state maps

The flexible factor analysis modeling used by mc-ASTRA allows known information to be incorporated at different levels of a tissue-state map while preserving variation learned from the data. In particular, sample-level covariates can be used to account for known sources of variation^35,36^, whereas feature-level priors can guide individual multicellular programs toward predefined molecular processes^34^. Because the resulting factors remain interpretable and compatible with downstream analysis frameworks, the same tissue-state maps can subsequently be used to characterize continuous patterns of tissue variation. We illustrate these capabilities in three applications spanning single-cell datasets of distinct biological contexts and experimental designs.

#### 2.2.1 Sample-level guidance can isolate known experimental variation while preserving reproducible disease-associated structure

We first tested whether a known source of sample variation could be explicitly guided in the tissue-state map while retaining disease-associated multicellular signals. We applied mc-ASTRA to ileum samples from the scIBD dataset ^44,45^, which included Crohn’s disease and control patient tissue samples collected across distinct tissue-enrichment fractions (i.e. the tissue compartment preferentially enriched during biopsy processing) and profiled using two library preparation protocols (Figure 2A). Because the enrichment fractions target different tissue compartments, we expected them to introduce systematic differences between tissues that could obscure disease-associated variation. Using samples of one chemistry as a reference dataset, we fitted a multicellular factor analysis model with the original author’s cell-type annotations, where one factor was guided using the categorical sample annotation of tissue-enrichment fraction. The guided factor showed the strongest association with tissue-enrichment fraction and explained a mean R² of 0.10 across cell types (ANOVA, adj. p-value = 1.65 × 10⁻⁹; Figure 2B–C), while Factor 2 was associated with Crohn’s disease status (ANOVA, adj. p-value = 0.002; mean R² across cell types of 0.08; Figure 2D) and reflected active inflammatory activity via JAK-STAT in enterocytes, goblet cells, and cycling transit amplifying cells (PROGENy activity, adj. p-value < 0.05). To test if the multicellular differences between control and disease tissues were reproducible independently of the library preparation, we projected the samples generated with the second protocol into the tissue-state map learned from the reference samples. Factor 2 scores remained associated with Crohn’s disease status in the projected samples (ANOVA, adj. p-value = 0.0004; Figure 2D), showing that guiding a known sample-level source of variation can concentrate enrichment-associated effects within the tissue-state map while preserving a disease-associated multicellular signal that is reproducible across library preparation protocols.

**Figure 2.**
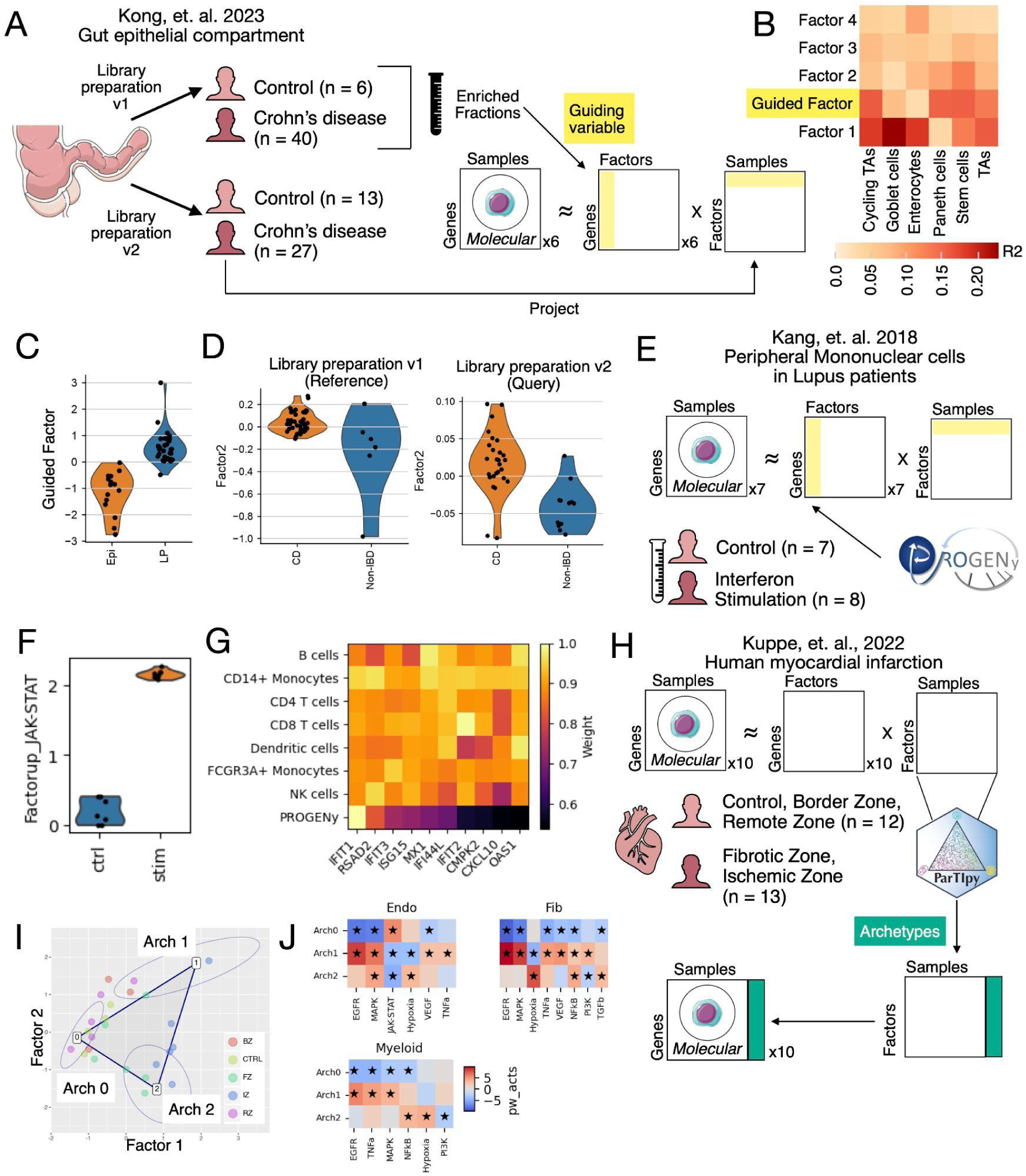
mc-ASTRA extends multicellular factor analysis with structured priors and downstream analyses. **A.** The epithelial compartment of the ileum of healthy and Crohn’s disease patients was analysed with multicellular factor analysis controlling for the technical effect of the enriched tissue compartment during biopsy treatment (i.e. enriched fraction). The model was fitted to a reference dataset containing a single library protocol (v1). The labels of the enriched fractions were used as structured priors to supervise the scores of a single factor, while letting the other factors recover the rest of the variance. A query dataset was projected afterwards to the inferred latent space. **B.** Proportion of explained gene expression variance of each cell-type and factor. **C.** Distribution of the scores of the guided factor for samples enriched for lamina propia (LP) or epithelium (Epi) **D.** Distribution of the scores of the factor associated with disease grouped by disease class for the reference and query dataset. **E.** Interferon-stimulated and unstimulated peripheral blood mononuclear cells of Lupus patients were analysed with multicellular factor analysis with a factor guided by the responsive genes to JAK-STAT as defined by PROGENy. **F.** Distribution of the factor guided by responsive genes to JAK-STAT activation grouped by stimulated and unstimulated samples. **G.** Gene weights (loadings) of the top 10 most responsive genes to JAK-STAT based on PROGENy, as well as their original weights. All weights are normalized for a maximum value of 1. **H.** Heart tissue samples of control and myocardial infarction patients across distinct regions and time-points were analysed with multicellular factor analysis. The generated latent space then was the input for archetypal analysis with ParTIpy that allows to identify extreme points that describe the variability of measured patients. The molecular profile of these archetypes can then be reconstructed using the model weights. **I.** Distribution of patient samples across the two factors associated with disease labels together with the location of three archetypes describing the variability of all samples. **J.** Pathway activities characterizing the estimated archetypes for three major cell-types. Stars reflect adjusted p-value of enrichment < 0.05. Colon and cell images provided by Servier Medical Art (https://smart.servier.com), licensed under CC BY 4.0.

#### 2.2.2 Feature-level supervision can contextualize molecular prior knowledge across cell types

Existing pathway signatures provide interpretable summaries of known signaling responses, but their gene weights are typically derived across biological systems and may not capture how the same pathway is expressed in individual cell types in a new multicellular context. We therefore asked whether mc-ASTRA could use a known perturbation-response signature as an anchor for a multicellular program while allowing the contribution of individual genes to be learned from the observed system, similarly as how it has been presented by the tool *Spectra^46^*. In contrast to applying a fixed pathway score, this analysis tests whether prior knowledge can identify the biological process of interest while mc-ASTRA contextualizes its transcriptional response across cell types.

To test this capability we analyzed peripheral blood mononuclear cells from patients with lupus before and after *ex-vivo* interferon-β stimulation^47^. Interferon-β is a type I interferon that signals through the Janus kinase/signal transducer and activator of transcription (JAK-STAT) pathway to trigger antiviral, antiproliferative, and immunomodulatory effects^48^. PROGENy^49^ provides a set of JAK–STAT-responsive genes whose weights summarize responses across multiple biological contexts. Thus, given available experimental data of a given perturbation in a multicellular system and prior knowledge of responsive genes, our objective was to build a multicellular factor analysis model focused in JAK-STAT responses, where it was possible to extract contextualized cell-type specific responses (Figure 2E). We fitted a two-factor model with non-negative factors and feature weights in which one of the factors was guided with all the genes whose expression increased upon JAK-STAT stimulation based on the top 500 most responsive genes reported in PROGENy (Methods). The guided factor explained, on average, 23% of the variance recovered by the model across cell types and was associated with interferon stimulation (ANOVA, adj. p-value = 7.88 × 10⁻¹³; Figure 2F), while remaining to be enriched by genes of the original signature (Methods; enrichment analysis, adj. p-value < 0.05). The second factor explained 75% of the recovered variance and was also associated with stimulation, but instead captured genes with lower activity after treatment (Methods; enrichment analysis, adj. p-value < 0.05). Because the model learns separate feature weights for each cell-type and factor, we could extract cell-type-specific versions of the original pathway signature. Among the ten most positively responsive genes based on PROGENy, we observed little correspondence between the original PROGENy weights and the reweighted signature (Pearson correlation = 0.07; Figure 2G). Thus, the prior anchored the factor to a known JAK–STAT response without fixing the contribution of individual genes, allowing the model to adapt the pathway signature to the cellular context represented in the data.

#### 2.2.3 Identifying archetypal tissue-states that describe continuous disease variation

Patients with the same disease rarely follow a single molecular trajectory. Instead, their tissues can show different combinations of remodeling processes. mc-ASTRA tissue-state maps can be used to describe this variation using archetypal analysis^50^, which focuses on the boundaries of the observed tissue-state map. Archetypal analysis identifies characteristic extreme states, or archetypes, that define these boundaries and then represents each sample according to its position relative to these extremes. Samples within the map can be interpreted as combinations of the processes represented by the archetypes, rather than being assigned to discrete groups. This is particularly useful for continuous and heterogeneous disease processes, for which discrete patient groupings may provide an incomplete description of the underlying biology. To illustrate this approach, we used ParTIpy^43^ to identify tissue archetypes in a mc-ASTRA map of myocardial infarction samples. We analyzed a single-nucleus dataset comprising healthy and infarcted human heart tissues sampled across different anatomical regions and stages of post-infarction remodeling spanning variation in major remodeling processes, including immune infiltration, cardiomyocyte stress, and scar formation^51^ (Figure 2H). The dataset contained three major groups: myogenic samples that reflected intact cardiac muscle from control patients or from remote and border zones regions of myocardial infarction patients; ischemic zone samples that represented the most damaged regions upon myocardial infarction, and fibrotic zone samples that captured the scarring processes upon cardiac damage.

After fitting a five-factor model, two factors associated with tissue class were used to define the latent space for archetypal analysis (ANOVA, adjusted P < 0.05; mean R² across cell types = 0.24). We identified three archetypes, which provided a stable representation of the data (Methods). Control, remote, and border-region-like samples were concentrated near a single archetype (Arch 0), potentially reflecting a healthy tissue state, whereas ischemic and fibrotic samples occupied continuous positions toward two injury-associated extremes (Arch 1 and 2; Figure 2I). Because each archetype has coordinates in the mc-ASTRA tissue-state map, we reconstructed its multicellular expression profile from the corresponding factor weights and estimated pathway activities from these profiles. One injury-associated archetype (Arch2) was characterized by expected inflammatory and fibrotic responses upon myocardial infarction, whereas the other (Arch1) showed MAPK, EGFR, and VEGF signaling, consistent with distinct stress-response and angiogenic processes reported previously^52^ (Figure 2J), showing how rather than assigning samples to these biological functions, archetypal analysis represented each sample according to its position between them. This application is an example of how the interoperability of mc-ASTRA tissue-state maps with downstream analytical frameworks can facilitate the interpretation of continuous combinations of multicellular remodeling programs that may be obscured by discrete sample stratification.

### 2.3 mc-ASTRA nominates spatiotemporal multicellular programs of gut healing

As a final application of mc-ASTRA as an integrated framework we characterized spatiotemporal tissue remodeling processes of gut damage and recovery. We analyzed a longitudinal MERFISH spatial single-cell atlas with a 940-gene panel of the distal colon from a dextran sodium sulfate (DSS)-induced mouse colitis model^53^. The dataset comprised samples collected before treatment (day 0), during early damage (day 3), at peak inflammation (day 9), and after DSS withdrawal and recovery (day 35) (Figure 3A). DSS treatment is expected to progressively disrupt epithelial integrity and induce inflammatory infiltration in the distal colon. Shortly after DSS withdrawal, tissue damage and inflammation can continue to increase before transitioning into a recovery phase characterized by epithelial regeneration, resolution of inflammation, and longer-term stromal and structural remodeling^54^. These coordinated molecular, compositional, and spatial changes provide an opportunity to use mc-ASTRA to represent tissue-state variation across the course of damage and recovery and resolve the multicellular programs associated with these transitions.

**Figure 3.**
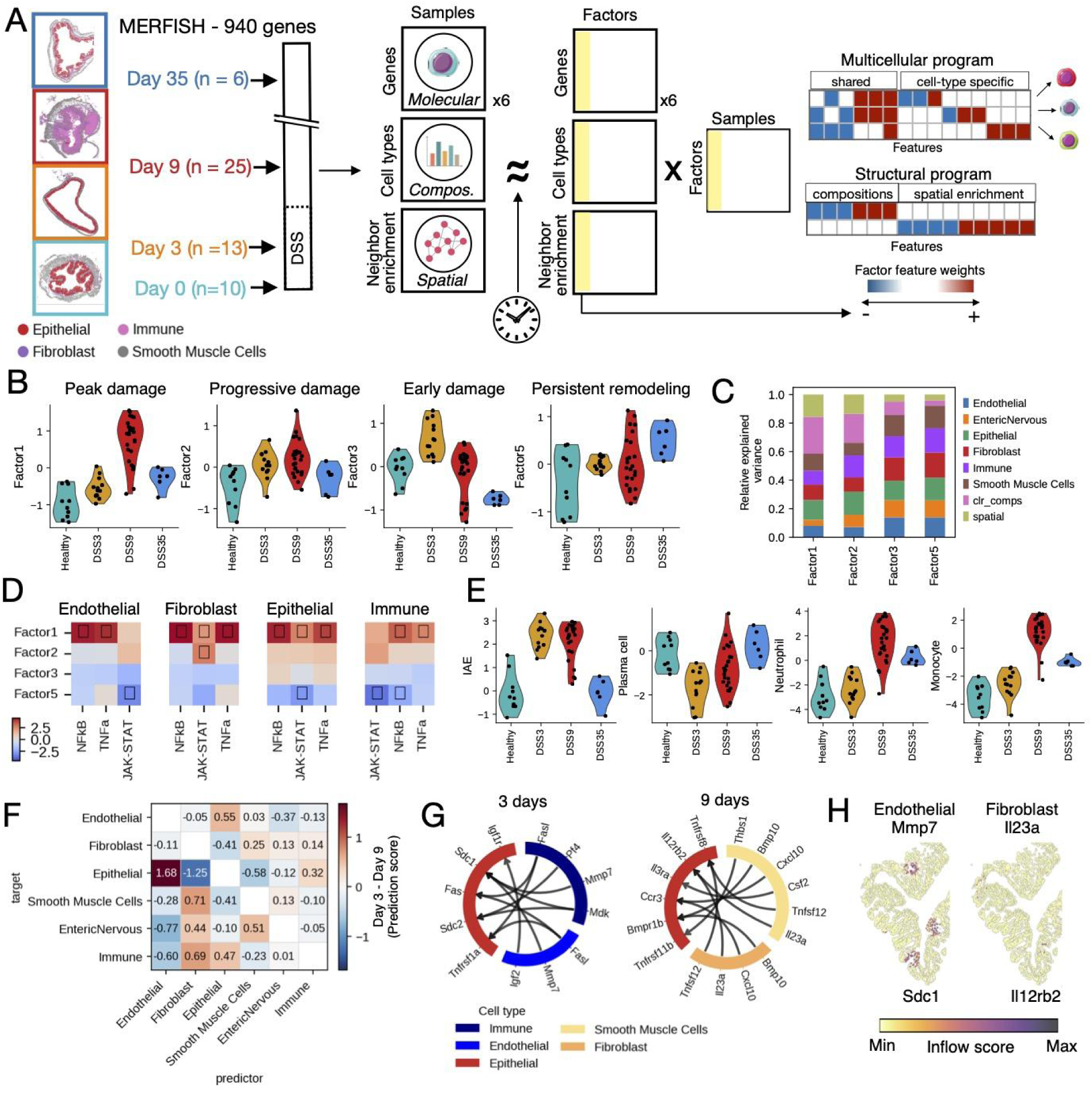
Spatiotemporal multicellular programs of gut damage and healing. **A.** Each sample represents tissue collected at a specific time point relative to dextran sodium sulfate (DSS) challenge. Arrows indicate the sampling time points. Example MERFISH images from each time point are shown below the time course and framed according to the corresponding time-point color. Four major cell types are annotated (Left). Graphical description of the multicellular factor analysis fitted with mc-ASTRA and the derivation of multicellular programs (Right). **B.** Factor scores of the four selected factors representing peak, progressive, and early damage, and persistent remodeling. **C.** Relative proportion of variance of each view explained by each factor. **D.** Signaling pathway activities inferred with PROGENy from factors’ gene loadings. Activities are highlighted with a small square if the enrichment adj. p-value was lower than 0.05 **E.** Centered-log-ratio transformed proportions of cell-states across time points. IAE refers to inflammation associated epithelial cells **F.** Differential multicellular coordination between day 3 and day 9 calculated as the contrast of multicellular information networks (See Methods). **G.** Top 10 ligand receptor interactions where epithelial cells were the receiver cells and immune or endothelial, and fibroblasts or smooth muscle cells were the sender cell types, for early (day 3) and peak (day 9) inflammatory programs respectively inferred from model factor weights. **H.** Inflow score of signals received by epithelial cells via Sdc1 or Il12rb2 receptors and Mmp7 and Il23a expressed by endothelial cells and fibroblasts, respectively.

First, we used mc-ASTRA to generate sample-level representations capturing molecular, compositional, and spatial tissue descriptors from the original MERFISH data (Methods). The compositional representation contained centered log-ratio-transformed cell-state proportions for each sample. The spatial representation quantified the pairwise proximity of cell states within each tissue. The molecular representation used a multiview structure containing pseudobulk gene-expression profiles for each major cell type after quality control. We then compared the tissue-state maps generated from individual tissue descriptors, and GloScope, an alternative sample-mapping approach, to ask whether differences between disease stages were retained despite differences in the tissue features underlying the maps (Supplementary Results). All tissue-state maps separated samples by time point (Supplementary Figure 1), as shown by the high accuracy with which each sample’s time point could be predicted from its neighbors in the map (minimum F1 score = 0.93). These results suggest that multiple sample maps can potentially share comparable predictive capabilities despite incorporating distinct tissue descriptors. Thus, we focused subsequent analyses on a joint factor model fitted on all tissue descriptors, which allowed us to determine how molecular, compositional, and spatial changes jointly contributed to the trajectories of tissue damage and recovery. The integrated model, besides incorporating molecular, compositional, and spatial tissue descriptors simultaneously, used the time-course information to account for temporal dependencies between samples emulating the strategy presented in MEFISTO^35^.

We inspected the factors and their feature loadings of the integrated model to nominate disease-associated multicellular programs (Figure 3A). We identified four multicellular programs representing distinct but simultaneous trajectories of tissue remodeling. These included the expected patterns of damage peaking at day 9 and recovery at day 35 (Factors 1 and 2), early damage at day 3 (Factor 3), and persistent tissue remodeling processes that did not resolve as time progressed (Factor 5) (ANOVA, adj. p-value < 0.05; Figure 3B). Comparison of the proportion of variance in each tissue descriptor explained by these multicellular programs revealed that early and persistent processes mainly reflected molecular changes in the cells rather than structural tissue changes in cellular composition or spatial organization (Figure 3C). In addition, the top spatial loadings across factors were consistent with a progression from early, predominantly epithelial reorganization to broader disruption of epithelial–stromal architecture at peak damage, accompanied by progressively increasing immune–stromal organization and persistent remodeling of stromal–muscular compartments (Supplementary Figure 2A). These results show how mc-ASTRA facilitates the identification of temporal disease processes with distinct multicellular characteristics from factor analysis models fitted to multimodal and temporal data.

We next compared the two multicellular programs capturing distinct trajectories of damage after DSS treatment and subsequent recovery at day 35 (Factor 1 and Factor 2) (Figure 3B). Factor 1, which captured the largest fraction of variance across tissue descriptors (mean R² across views = 0.24), was activated only during peak inflammation at day 9 and resolved by day 35, with no activation at day 3. This peak inflammation program showed high activity of inflammatory signaling pathways such as TNFα, NFkB, and JAK-STAT across multiple cell types (Figure 3D), as well as expected increased presence of neutrophils and monocytes^55^ (Figure 3E). In contrast, in the progressive secondary program captured by Factor 2 (mean R² across views = 0.05), samples at day 3 already showed a damage state similar to that observed during peak inflammation at day 9, and was characterized by the early presence of inflammatory associated epithelial cells in the tissue and a reduction on plasma cells based on its factor feature loadings that were confirmed by the sample’s cell proportions (Figure 3E). Given the central role of JAK-STAT in integrating multiple inflammatory cytokine signals and its established therapeutic relevance in IBD through JAK inhibitors^56^, we compared the factor loadings for JAK-STAT-responsive genes in epithelial cells across the two programs. The gene weights of these genes pointed to distinct temporal expression patterns (Supplementary Figure 2B). For example, *Il15* upregulation and *Socs3* downregulation were more progressive, whereas *Isg15* and *Cxcl10* showed more abrupt increases (Supplementary Figure 2C). Comparing JAK-STAT-responsive gene weights across cell types also revealed cell-type-specific differences in *Ifit2* expression. In epithelial and immune cells we observed *Ifit2* to increase its expression gradually, while in fibroblasts and endothelial cells we observed a progressive downregulation, suggesting potential differential functions of this interferon responsive gene (Supplementary Figure 2B). These distinct gene regulatory patterns recovered by comparing the contribution of genes to the peak and progressive inflammatory programs, show how mc-ASTRA can be used to explore coordinated tissue remodeling mechanisms at distinct scales, from general multicellular responses and structural changes to specific gene responses within a cell type population.

We then focused on the early response program (Factor 3) to better characterise the starting inflammatory processes. In epithelial cells, among the top 15 genes with the highest overexpression loadings we observed *Fos*, *Fosb* and *Egr1* illustrating a MAPK/ERK-driven damage-response signaling and initiation of downstream programs for migration, cytokine production, and growth-factor signaling^57^ (Supplementary Figure 2D). In addition, we observed induction of *Mmp15* related to ECM remodeling required for cell migration and repair ^58^. Within the immune cell-types, we captured the overexpression of *Ccl22*, which may point to early regulatory T cell retention and disease suppression^59^ and the expression of *Il17re*, which could point to an “early-warning” cytokine circuit ^60^ (Supplementary Figure 2D).

To contrast the early and peak-damage responses, we calculated differential multicellular coordination between Factors 3 and 1 (Figure 3F; see Methods). Briefly, we quantified how well the molecular state of each cell type predicted that of every other cell type during early (Factor 3, day 3) and peak damage (Factor 1, day 9) and then calculated the difference between their predictive importances. We assumed that this difference reflected how much more coordinated is the transcriptional response between two cell-types at early versus late damage. At early damage, epithelial states showed stronger coordination with immune and endothelial compartments, while at peak damage we observed an increased association of epithelial cells’ activity with fibroblasts and smooth-muscle cells (Figure 3F). Ligand–receptor inference further showed a marked change in the cellular sources and functions of epithelial communication between early and peak damage (Figure 3G). At early damage, immune- and endothelial-derived interactions included signals associated with epithelial injury and survival, such as *Fasl–Fas* and *Igf2–Igf1r^61,62^*, together with *Mmp7–Sdc1*, linking extracellular proteolysis to epithelial barrier remodelling^63^. By day 9, during peak damage, we observed an increased inflammatory and other remodelling-associated signals involving *Tnfsf12*, *Cxcl10*, *Csf2*, *Il23a*, *Thbs1*, and *Bmp10 ^64–66^*. Spatial mapping of the *Mmp7–Sdc1* and *Il23a-Il12rb2* ligand-receptor interaction to the epithelium of day 3 samples (Figure 3H; see Methods), provided a visual example of how exclusive communication events are then distributed in the tissue. Together, these results support a progression from epithelial stress associated primarily with immune and endothelial inputs during early damage to a peak-damage state in which inflammation becomes increasingly integrated into the stromal response, coupling epithelial states to activated fibroblast and smooth-muscle compartments undergoing tissue remodelling.

Finally, we examined the multicellular program that accumulated across DSS exposure and remained altered after recovery to identify molecular features of residual damage (Factor 5). Despite the near restoration of mucosal architecture by day 35, the colon retained a coordinated post-injury state. In fibroblast and immune compartments we observed persistent *Axin2* expression suggesting ongoing Wnt-dependent niche activity^67^. Epithelial *Gdf15/Nlrc3* and immune *Nod2* indicate continued adaptation to cellular stress and microbial exposure ^68–70^. Increased *Adamts12* expression and activated fibroblast features were consistent with continued matrix and stromal remodeling ^71^. Loss of *Per1* across stromal and immune compartments, together with reduced *Irf8* in immune cells, could relate to incomplete restoration of circadian and myeloid homeostatic programs^72,73^. These patterns indicate that structural recovery was not accompanied by complete molecular restoration, but instead left compartment-specific residual states across stromal, epithelial, and immune cells, consistent with the persistence of inflammation-associated fibroblasts and increased plasma cells reported at day 35 (Supplementary Figure 2F-H).

## 3. Discussion

The growing complexity of single-cell and spatial tissue datasets requires frameworks that connect population-level sample variation with interpretable multicellular processes related to tissue function. Here, we presented mc-ASTRA, a Python framework for building and interpreting tissue-state maps from single-cell, spatial, and complementary sample-level data. mc-ASTRA integrates tissue descriptor generation, flexible multicellular factor analysis, and downstream analysis of multicellular coordination in a single workflow, enabling researchers to identify the technical and biological sources of variation across samples and relate them to coordinated molecular, compositional, and spatial tissue remodeling.

Across distinct extensively documented applications (https://saezlab.github.io/mc-astra/tutorials/), we illustrated the extended capabilities of multicellular factor analysis facilitated by the new classes of semi-supervised factor models used within mc-ASTRA. On the one hand, modeling guidance based on sample covariates allows expected technical and biological signals to be explicitly accounted for, while preserving latent variation beyond these annotations that may represent previously uncharacterized biological processes^35,74^. On the other hand, guidance using known biological groupings of features (e.g. pathway gene sets) allows to target specific multicellular processes, while adapting the prior knowledge to the specific biological context of interest^34^. This contextualization is relevant as most of the available pathway annotations and perturbation signatures are built from different experimental systems and disease signatures may be obscured by technical issues related to tissue handling^75^, among others. Together, these analyses show how mc-ASTRA can be tailored to different biological assumptions and experimental designs while supporting biological interpretation across scales, from general comparisons of tissue descriptors to specific hypotheses about cellular communication and cell-type-specific molecular processes.

Available evaluation frameworks for tissue-state maps often assess how well known annotations of tissue samples can be recovered^25^, providing useful metrics of information retention. However, this makes predefined variables part of the evaluation target and may overlook biologically structured variation beyond them. mc-ASTRA complements these approaches with reconstruction-based metrics that quantify the variance captured by the full model and individual factors, and relate this variance to tissue descriptors and covariates. This combination of metrics provides an extended assessment of sample representations by allowing users to determine not only which known variables are retained, but also which tissue components contribute to sample variability and how much of this variability is captured by annotated versus latent sources of variation.

While existing sample-representation frameworks largely focus on describing relationships between samples, mc-ASTRA uses the multicellular processes underlying these relationships as the starting point for downstream biological interpretation. Our analysis of gut damage and healing showed how spatiotemporal multicellular programs that involved distinct coordinated tissue activities could be studied simultaneously. We observed, for example, that the major progressive and peak inflammatory processes involved both the activation of specific molecular programs together with structural changes in the tissues. This contrasted with the observation that early and persistent damage involved mainly multicellular molecular reprogramming despite an almost intact mucosal architecture. The combination of mc-ASTRA’s compatibility with functional transcriptomics tools and its implementation of multicellular coordination network inference allowed us to contrast the potential intercellular signaling mechanisms behind early and peak damage. By connecting mc-ASTRA to cell-cell communication and spatial mapping tools^76,77^, we identified a rewiring of coordination relationships between cell-types, as well as differences in the ligand-receptor usage of epithelial cells. This analysis illustrates how multicellular programs identified from variation across tissues can provide an entry point for mechanistic interpretation, connecting coordinated tissue remodeling to candidate intercellular signaling mechanisms. More specialized network approaches are increasingly able to connect cell–cell communication with intracellular signaling, model multicellular responses to perturbations, or infer context-specific signaling networks from prior knowledge and omics data ^76,78,79^. We envision that integrating these approaches with the multicellular processes identified by mc-ASTRA could facilitate the prioritization of signaling effectors responsible for coordinated tissue remodeling.

The flexibility of mc-ASTRA gives users control over several aspects of the analysis. In single-cell spatial data, for example, tissue descriptors can depend on cell segmentation, cell-type annotation, the choice and parameterization of spatial metrics, and gene-expression filtering. In other settings, descriptors could incorporate additional omics modalities (e.g. proteomics and morphology^32^: https://saezlab.github.io/mc-astra/notebooks/CRC_morpho/) or be defined around spatial niches rather than individual cell types, as may be useful for spot-based technologies such as Visium^37^. These analytical choices can influence the resulting tissue-state maps and their biological interpretation and therefore need to be considered in the context of the biological question and data. Similar considerations apply to factor modeling, where the increased flexibility of the framework introduces choices among different priors, guiding variables, and model configurations. Through extensive documentation, we provide new users with a collection of template models for distinct applications and data combinations. However, these choices should still be evaluated within each biological context using appropriate model diagnostics.

In summary, mc-ASTRA provides a flexible framework to construct and interpret tissue-state maps from single-cell and spatial data, linking variation between individual samples to coordinated multicellular processes and tissue organization. We envision these representations as more than endpoints for molecular stratification. Their integration with circulating biomarkers, genetic variation, imaging, and longitudinal clinical information could connect mechanistic descriptions of tissue dysfunction with population-scale patterns of disease. At the same time, their mechanistic interpretability could provide a reference for evaluating and designing experimental and computational models according to the disease states they reproduce. More broadly, incrementally extensible and interpretable tissue-state maps could provide a shared scaffold through which molecular, clinical, and experimental descriptions of disease are progressively connected rather than studied in isolation.

## 4. Methods

### Data and code availability

mc-ASTRA is available as a Python package (https://github.com/saezlab/mc-ASTRA), the documentation has the code to reproduce the results section 2.2 (https://saezlab.github.io/mc-astra/tutorials/). Code to reproduce the analysis on the induced colitis dataset is available at: https://github.com/saezlab/mc-ASTRA_pub. All data used came from original sources listed in the methods section. A Zenodo folder with data needed to reproduce the results is available at: https://doi.org/10.5281/zenodo.22280030.

### multicellular Analysis of Sample Tissue Representations and Associations (mc-ASTRA)

mc-ASTRA is a Python package that provides tools needed to construct sample or tissue-state maps from tissue-level descriptors inferred from single-cell molecular data and other clinical or profiling technologies. As such, it bridges pre-processing and quality control metrics for single-cell -omics, with factor modeling strategies and mechanistic interpretation utilities to represent sample variability as a function of activation of multicellular processes. The core strategy of mc-ASTRA is to transform the data of a patient cohort into a multiview representation, where each view captures a specific tissue descriptor of the cohort, and to decompose it into an interpretable latent space of lower dimensions. This means that for a sample cohort profiled with single-cell technologies, mc-ASTRA reduces the information of thousands of cells and features, into a collection of latent programs (i.e. multicellular programs) that describe the observed variability of the tissues.

mc-ASTRA relies on probabilistic group factor analysis for the construction of the latent space, which is a strategy for simultaneous low-rank factorization of multiple data matrices. Generally, group factor analysis decomposes multiple observation matrices (i.e. views) across different batches (i.e. groups) as the product of a factor score matrix with K components and a collection of weight matrices that summarize the contribution of each feature in each view for each factor k ∈ K. Group factor analysis provides three major outputs relevant for the construction of tissue-state maps: i) a latent space represented by the factor scores per sample, ii) the contribution of each feature and tissue descriptor to each latent variable (i.e. feature loadings), and iii) reconstruction errors per latent variable that allows to attribute its importance in describing tissue descriptors (i.e. coefficient of determination, R²). Downstream analyses take advantage of the feature loadings of each factor, that are interpreted as multicellular programs that capture coordinated activities across cell types, together with other remodeling features, such as compositional or spatial organization. We refer to this complete approach as multicellular factor analysis.

mc-ASTRA relies on the probabilistic implementation of factor analysis provided by MOFA-FLEX ^33^ that allows for the definition of models with flexible priors and learning structures, including supervision signals at the level of features or samples together with group-aware decomposition. These novel characteristics are on top of the model’s ability to handle missing values at different levels. mc-ASTRA’s objective, thus, is to facilitate the usage of these complex models in the construction of patient or sample maps, latent spaces whose coordinates define specific combinations of tissue or patient descriptors.

### Building multiview representations of tissues or patients

For standard multicellular factor analysis for single-cell molecular data, mc-ASTRA takes as input an *anndata* object with cell annotations and creates a multiview representation of patients where each view corresponds to the pseudobulk expression profile of a cell-type. We used *decoupler-py*’s ^80^ pseudobulk representations, together with its filters for lowly-expressed features to define cell-type views. At the level of patients, it is possible to filter samples within a view based on the number of cells used to build the pseudobulk and the number of features, and across the multiview representation based on the number of non-empty views. Within views, features can be excluded based on prior knowledge, for example known contamc-ASTRAtion patterns. *Scanpy*’s ^81^ log-normalization and highly-variable feature selection is used within pre-processing functions adapted for multi-view representations.

mc-ASTRA facilitates the inclusion of compositional and spatial views, representing cell-type/state proportions and spatial organization between cells or features. For compositional views, centered-log-ratios (clr) are used as standard, while for spatial views spatial neighborhood enrichment from *squidpy* is ported ^77^. For guided factor models, mc-ASTRA interfaces between prior knowledge databases and enrichment tools via decoupler-py to build the data structures with feature annotations needed by MOFA-FLEX, reducing the friction between feature set selection and model definition.

### Model representation and downstream analyses

To facilitate the compatibility of factor models with visualization and analysis tools available for anndata objects, we provided an anndata data structure that collects MOFA-FLEX models and the data used to train them. This allows users to natively use, for example, Scanpy’s clustering, embedding, and visualization functions to define and visualize patient groups or *ParTIpy*’s archetypal analysis toolbox to identify the most characteristic samples in the map ^43^. In addition, we have added to this toolbox parametric statistical tests for associations of factors with continuous or categorical annotations of the samples with an optional use of random effects in mixed-effect models. For enrichment tools we wrapped decoupler-py’s univariate linear model to work in a multi-view setup that allows to calculate enrichment scores for each view individually.

### Reconstruction of multicellular dependency networks and cell-cell communication

For each latent factor of interest, we reconstructed multicellular networks from multicellular molecular views that represent how much of the activity of a cell type can be predicted by the activity of others, turning multicellular coordination inference from an unsupervised task to a predictive task. Feature loadings were first partitioned by data view and direction (positive and negative), with each view corresponding to a cell type or other molecular compartment represented in the model. Within each selected view, features with the strongest positive and negative loadings for the factor were identified using a user-defined percentile threshold. These gene sets were then scored in the corresponding sample-level original representation (e.g. pseudobulk expression profiles for cell types) using the univariate linear model enrichment method implemented in *decoupler-py* ^80^. Missing expression values were replaced with zero before enrichment. The resulting activity scores were concatenated across views, producing one sample-by-view matrix for positively associated programs and another for negatively associated programs.

Directed multicellular networks were inferred independently from the positive and negative activity matrices. For each view in turn, its program activity was treated as the response variable, while the activities of all remaining views were included simultaneously as predictors in a multivariable linear model. When a grouping variable was supplied, a linear mixed-effects model with a group-specific random intercept was fitted instead, thereby accounting for repeated measurements or other clustered sample structures. Features can optionally be standardized to zero mean and unit variance before model fitting, and samples containing missing values were removed jointly across all variables included in a model.

Each predictor–target combination defined a directed network edge. The edge coefficient represented the conditional association between the predictor activity and the target activity after adjustment for all other views. Overall model fit was quantified using the R² of the multivariate models. For mixed-effects models, marginal (R²) was calculated as the proportion of total variance attributable to the fixed effects, with total variance defined as the sum of the fixed-effect, random-intercept, and residual variances. An edge score was subsequently calculated as the regression coefficient multiplied by the corresponding model-level (R²) with the objective of representing coordination as effective prediction and strong effect.

Potential cell-cell communication events constrained by these multicellular networks can be extracted by identifying coherent ligand-receptor co-expression patterns. To define coherent ligand-receptor interactions, we first defined all possible pairs of communicating cell-types using the edge scores of the multicellular networks. By default, only positive interactions representing coordinated upregulation of the multicellular programs are kept. Then, based on the sign of the partitioned features of the factor of interest (i.e. positive or negative), ligand-receptor interactions are selected based on coherent signs and the value of the multiplication of their factor loadings. Ligand-receptor interactions are extracted from LIANA+ consensus database ^76^.

### Multicellular programs of gut damage in Crohn’s disease patients

The epithelial compartment of the patient cohort profiled in Kong, et. al. 2023^45^ was obtained from the scIBD dataset available at the Broad Single Cell Portal (SCP2927)^44^. Original cell-type annotations were used to build the molecular views for multicellular factor analysis, however samples were separated into reference and query subsets based on their library preparation protocol (v1 = reference, v2 = query). Molecular sample-level representations were built as a multi-view representation of pseudobulk expression profiles, with views defined by cell type. We retained pseudobulk profiles calculated from at least 20 cells and cell-type views represented in at least 40% of all samples. Within each view, we kept genes with a minimum expression of 20 counts in at least 40% of the available samples and excluded views containing fewer than 100 genes after filtering. Samples whose expression profiles covered less than 90% of the feature space of a view were removed, after which the minimum 40% sample representation criterion was applied again. Expression profiles were log-normalized to a total count of 1,000,000 per sample and centered. Highly variable genes were subsequently retained for each view using scanpy’s “Seurat” flavor^81,82^. Finally, samples represented in fewer than five views were excluded from the multi-view representation. We fitted a MOFA-FLEX model with 5 factors to the reference data with a horseshoe weight prior and a guiding variable assigned to a factor using a Bernoulli likelihood. We used a learning rate of 0.01. To project the query dataset we multiplied the pseudobulked gene expression of each cell-type with the pseudoinverse of the feature loadings of the model.

### Contextualized JAK-STAT responsive genes in lupus

Demultiplexed scRNA-seq count matrices of peripheral mononuclear cells from eight pooled lupus patients samples, each before and after IFN-beta stimulation were downloaded using LIANA+ ^47,76^. Molecular sample-level representations were built as a multi-view representation of pseudobulk expression profiles, with views defined by cell type. We retained pseudobulk profiles calculated from at least 10 cells and cell-type views represented in at least 40% of all samples. Within each view, we kept genes with a minimum expression of five counts in at least 40% of the available samples and excluded views containing fewer than 100 genes after filtering. Samples whose expression profiles covered less than 90% of the feature space of a view were removed. Raw counts were stored before normalization, and expression profiles were log-normalized to a total count of 1,000,000 per sample and centered. Finally, samples represented in fewer than five views were excluded from the multi-view representation. For each view, we created an annotation of all genes whose expression increases upon JAK-STAT stimulation based on PROGENy’s top 500 genes. A two-factor model was fitted with MOFA-FLEX, using an informed horseshoe prior with a 75% annotation confidence that captured in a single factor the contextualized response to JAK-STAT based on the data. Non-negativity of scores and feature weights was enforced to allow guidance and we used a learning rate of 0.001. Two factors were chosen because of sample size and because it could facilitate the interpretation of genes as up and downregulation upon perturbation. To test if the created factors associated with the original JAK-STAT responsive genes, we calculated pathway activities with PROGENy, using decoupler-py’s univariate linear model.

### Archetypical myocardial infarction patients

Single nuclei RNA-seq (sn-RNA-seq) gene count expression matrices from 27 human heart tissue samples encompassing healthy and myocardial infarction patients from Kuppe, et. al. 2022 were downloaded from cellxgene (https://cellxgene.cziscience.com/collections/8191c283-0816-424b-9b61-c3e1d6258a77)^51^. Original cell-type annotations and disease labels were used during the analysis. Molecular sample-level representations were built as a multi-view representation of pseudobulk expression profiles, with views defined by cell type. We retained pseudobulk profiles calculated from at least 10 cells and cell-type views represented in at least 40% of all samples. Within each view, we kept genes with a minimum expression of 20 counts in at least 40% of the available samples and excluded views containing fewer than 100 genes after filtering. Samples whose expression profiles covered less than 95% of the feature space of a view were removed. Expression profiles were log-normalized to a total count of 1,000,000 per sample and centered. Highly variable genes were subsequently retained within each view using scanpy’s “Seurat” flavor ^81,82^. Finally, samples represented in fewer than five views were excluded from the multi-view representation. A 5-factor model was fitted using MOFA-FLEX with a horseshoe weight prior and a learning rate of 0.001. Two factors associated with the disease labels were identified with an ANOVA (adj. P-value < 0.05) and further used for archetypal analysis using ParTIpy^43^. Based on low values of information criteria and bootstrap variance, and high proportion of explained variance, we selected three archetypes. The gene expression profile of these three archetypes was reconstructed by multiplying the factor scores with the gene weights of the two selected factors.

### Spatiotemporal multicellular programs of gut damage and healing

MERFISH data from Cadinu, et. al. was downloaded from https://doi.org/10.5061/dryad.rjdfn2zh3. The two single-cell objects containing the data from day 3 to day 21 and the data from day 35 were merged into a single object. Tier 1 original author annotations were used for all molecular sample representations, since they contained the eight major compartments: epithelia, fibroblast, immune, endothelial, smooth muscle, neurons and glial, interstitial cells of Cajal, and adipose. Tier 2 and 3 original author annotations were mapped into a new annotation (Tier mc-ASTRA) of 26 cell-states for compositional and spatial tissue descriptors (Supplemental Table 1). Marker genes of Tier 1 were calculated by contrasting the pseudobulk expression profiles of one cell-type versus the rest using Wilcoxon tests. Pseudobulk profiles were log normalized with a scale factor of 10,000 before performing the tests. We selected as markers all upregulated genes with a log fold change equal or greater than 1.5 and an adjusted Benjamini-Hochberg p-value less than 0.05.

Molecular sample-level representations were built as a multi-view representation of the pseudobulks of the raw counts of all cells grouped by their Tier 1 annotation. We kept all profiles whose pseudobulk expression was calculated from 10 or more cells and kept all cell-types that had profiles of 90% of all samples and at least 50 genes. For each view we kept genes whose minimum expression of five counts was observed in at least 40% of all samples within the view. In addition, we filtered out all samples whose expression profile covered less than 95% of the feature space. Exclusive marker genes of each cell-type were considered contaminants of the rest and excluded from their views if present. Before model fitting, data was log-normalized with a size factor of 100,000. Day 21 samples had a mean Pearson correlation across samples and cell-types of 0.04, reflecting a high technical effect and thus were excluded for this analysis. The compositional sample-level representation contained centered-log-ratio proportions of each cell-state in Tier mc-ASTRA for each sample. The spatial sample-level representation contained neighborhood enrichment scores of each pair of cell-states in Tier mc-ASTRA for each sample as implemented in *squidpy^77^*. We used 1000 permutations of the spatial graph and kept the interactions between the same cell-states.

We fitted four factor models with MOFA-FLEX^33^: Three of them used each sample-representation individually and the fourth one combined all the sample-representations and included a factor prior of the time course using gaussian processes, as presented in MEFISTO^35^, but with a Horseshoe weight prior. All models were fitted to recover 10 factors, with a learning rate of 0.001. Models fitted with individual sample-representations used a horseshoe prior for the feature weights. In addition to the sample maps generated with mc-ASTRA, we used *GloScope^8^*to calculate a pairwise distance matrix between samples with *patpy’*s*^25^*gpu implementation and the principal component analysis latent space of single cells calculated by the original authors.

Information retention of the time points of each sample map was evaluated using *patpy*. The objective of this evaluation was to test how accurate is the prediction of the time point of a sample based on its immediate neighbors. We used the five nearest neighbors for the classification and reported macro-calibrated F1 scores. To calculate the proportion of feature variance associated with the time course for each of mc-ASTRA’s sample maps, we first tested for association of individual factors to the time course labels (adjusted p-value < 0.05) and then for associated factors we summed their corresponding proportion of explained variance per view. Associations were tested using analysis of variance (ANOVA) and p-values were corrected using the Benjamini-Hochberg procedure.

To interpret the model that included all tissue descriptors we first ensured that for all selected factors the activation of the multicellular program was positive relative to control samples (day 0). This also affected their corresponding feature weights. Relative explained variance per view and factor was calculated by making the original R² sum to 1 for each factor. Pathway activities per factor and cell-type were calculated using decoupler-py^80^ and PROGENy’s^49^ top 500 responsive genes. Spatial mapping of ligand-receptor interactions was done using the inflow score of LIANA+^76^ using default settings and a bandwidth of 50.

Multicellular dependency networks were calculated from the 0.8 percentile of the positive loadings of Factor 1 and Factor 3. To identify differences between cell dependencies stronger in early over late time points, the corrected dependency estimates were contrasted between Factor 3 and Factor 1.

## Use of AI

Commercial large language models were used for the editing of grammatical errors of this manuscript

## 5. Acknowledgements

RORF was supported by the European Research Council (ERC) Synergy Grant 101118531 awarded to JSR. Thanks to Leonie Kuchenhoff for providing the template code for compositional analysis and revising the text. We thank Ines Rivero, Daniele Bottazzi, Eduardo Villablanca and Martin Suarez for comments on the manuscript. We thank Ilia Kats, Florin Walter, and Arber Qoku for the technical support given during the implementation of the MOFA-FLEX models.

## 7. Authors Contributions

RORF: Conceptualization, Data curation, Software, Formal analysis, Validation, Investigation, Visualization, Supervision, Methodology, Writing – original draft. KASKS, FGV and LV: Software. JSR: Resources, Supervision, Funding acquisition, Writing – review and editing.

## 9. Supplementary Materials

### Supplementary Results

#### Tissue-state map comparison

We fitted four factor-analysis models with 10 factors each. Three models used the compositional, spatial, or molecular representation separately. A fourth model jointly incorporated all three representations and included a Gaussian-process prior based on the time course to model the trajectory of tissue damage and recovery (Methods). To compare to an alternative state-of-the-art sample map, we used GloScope^8^ to calculate pairwise distances between samples directly from the single-cell MERFISH data. Uniform Manifold Approximation and Projection (UMAP) was used to visualize in two dimensions the latent spaces from the four factor analysis models and the distance matrix generated with GloScope (Figure 2C). All maps separated samples by time point, as shown by the high accuracy with which each sample’s time point could be predicted from its neighbors in the map (Figure 2D). The sample map built with molecular data was the only one achieving a perfect classification performance (F1 = 1), while the one built using only the spatial representation performed the worst (F1 score = 0.93), followed by the one using compositional data (F1 score = 0.95). GloScope and the model incorporating all tissue descriptors had similar classification performance (F1 score = 0.97). However, although UMAP distances cannot be compared quantitatively across representations, GloScope visually placed day 35 samples (recovery) in an isolated region instead of reflecting its similarities to healthier time-points (day 0 and 3). These results suggest that multiple sample maps can potentially share comparable predictive capabilities despite incorporating distinct tissue descriptors. Thus, interpreting tissue remodeling requires examining which modalities and tissue features contribute to each inferred process, rather than evaluating sample separation alone.

To identify which tissue descriptors contributed most to variation among samples across time points, we quantified the proportion of variance explained in each view by each model. In the single-representation models, differences among time points accounted for 94% of the variance in cell-state composition and 58% of the variance in spatial organization, indicating that changes in cellular compositions are more associated with the disease stage than changes in spatial organization. In the multicellular molecular model, we quantified which cell-type molecular variability associates the most with disease stage. We observed that smooth muscle cells, fibroblasts, and epithelial cells were the top three cell-types whose transcriptional variability better associates with the disease trajectory (84%, 75%, and 78%, respectively; Figure 2E). The model that incorporated all tissue descriptors captured similar trends as the models based on individual representations, with the advantage of directly characterizing which tissue descriptor has coordinated changes with others (for example, which gene expression programs are activated under certain structural changes in the tissue). When comparing all factors in the full model associated with the disease stage (ANOVA, adj. p-value < 0.05), the major compositional and spatial organization changes were mostly associated with the molecular variability of epithelial cells (Figure 2F). In contrast, factors capturing greater variance in gene expression of fibroblasts immune and smooth muscle cells explained less variance of compositional and spatial features (e.g. Factor 2), suggesting the potential existence of molecular responses upon tissue damage that are independent of the tissue architecture. These results illustrate how mc-ASTRA complements sample-level classification evaluations of latent spaces by identifying the tissue descriptors and specific latent factors that may contribute to disease progression.

### Supplementary Tables

**Supplementary Table 1.** Taxonomy of tissue-state mapping methods. Frameworks for sample mapping from single-cell and spatial data apply a variety of representation learning techniques for multimodal data. Major differences between available methods are the flexibility of tissue representation inputs and native interpretability.

| Class | Subclass | Tools | Description |
| --- | --- | --- | --- |
| Distance-first distributional methods | N/A | GloScope<sup>8</sup>, PILOT<sup>10,11</sup>, QOT<sup>9</sup>,<br><sup>12</sup>scSLIDE | Represent each sample as a distribution of cells in a shared cell-state space, then produce pairwise-distances. The sample map is usually obtained by embedding the resulting sample-by-sample distance matrix and does not decompose variance across biological views. Thus no direct relationship exists between the embedding and single features. |
| Multimodal data reconstruction | Deep patient embedding methods | MrVI<sup>14</sup>, scPoli<sup>13</sup>, PaSCient<sup>16</sup>,<br>PULSAR<sup>17</sup>,<sup>15</sup>Phenoverse | Learn sample embeddings from raw single-cell data or integrated atlases using variational autoencoders, attention pooling, transformers, or hierarchical generative models. They often optimize integration, prediction, reference mapping, counterfactual analysis, or perturbation simulation. Latent dimensions are less directly interpretable, recovered variance is usually unavailable, and the use of multiple tissue descriptors is not standard. |
|  | Tensor and factor methods | scITD<sup>18</sup>, Tensor-cell2cell<sup>22,23</sup>,<br>MUSTARD<sup>21</sup>, scPAFA<sup>20</sup>,<br>MOFAcell<sup>19</sup>, **mc-ASTRA** | Embed sample-level multicellular data using interpretable factor analysis or tensor decomposition. For each model it is possible to extract reconstruction errors, and view and feature importances. **mc-ASTRA is the most general member because it allows heterogeneous views, missingness, structured priors, and patient-map interpretation in one workflow, overcoming fixed tensor assumptions.** |
|  | Compositional and pseudobulk baselines | scECODA<sup>24</sup>, pseudobulk,<br>cell-type pseudobulk | Rely on simpler representations and embedding strategies. Their main disadvantage is their lack of a multimodal focus. Their interpretation can be obscured by multiple tissue descriptors |

### Supplementary Figures

**Supplementary Figure 1.**
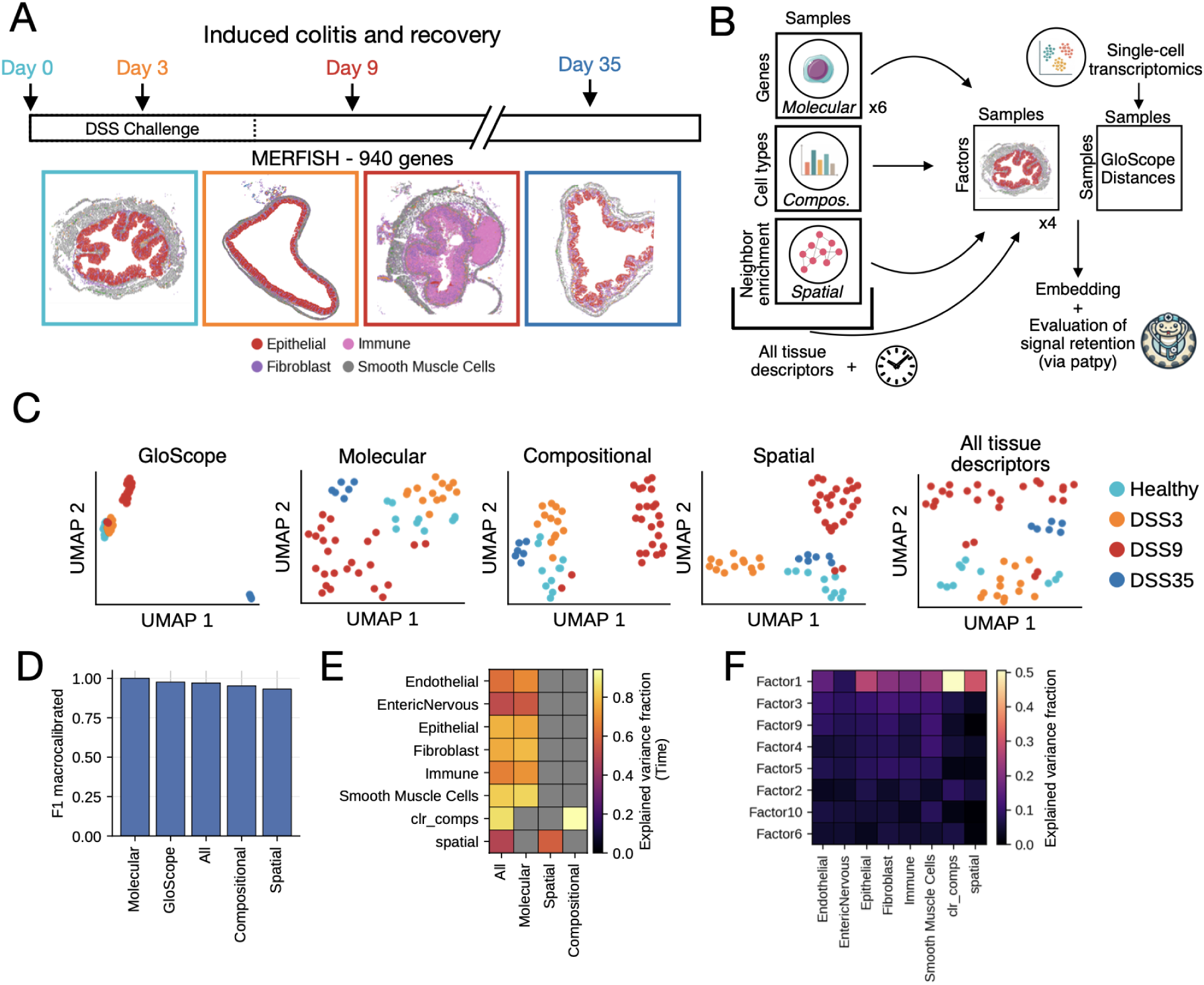
Estimating the importance of tissue descriptors in sample-level variability. **A.** Experimental design of the induced colitis and recovery dataset in mouse distal colon. Each sample represents tissue collected at a specific time point relative to dextran sodium sulfate (DSS) challenge. Arrows indicate the sampling time points. Example MERFISH images from each time point are shown below the time course and framed according to the corresponding time-point color. Four major cell types are annotated. **B.** Tissue descriptors and tissue-state maps generated with mc-ASTRA. Four mc-ASTRA models were fitted using molecular, compositional, or spatial descriptors individually, or all three descriptors together with time guidance. A GloScope tissue-state map was additionally generated using patpy. **C.** Uniform Manifold Approximation and Projections (UMAP) of the five tissue-state maps colored by time point. **D.** Evaluation of time-point information retention by the different tissue-state maps. **E.** Proportion of variance of each view explained by individual models. **F.** Proportion of variance of each view explained by each factor of the model that included all tissue descriptors.

**Supplementary Figure 2.**
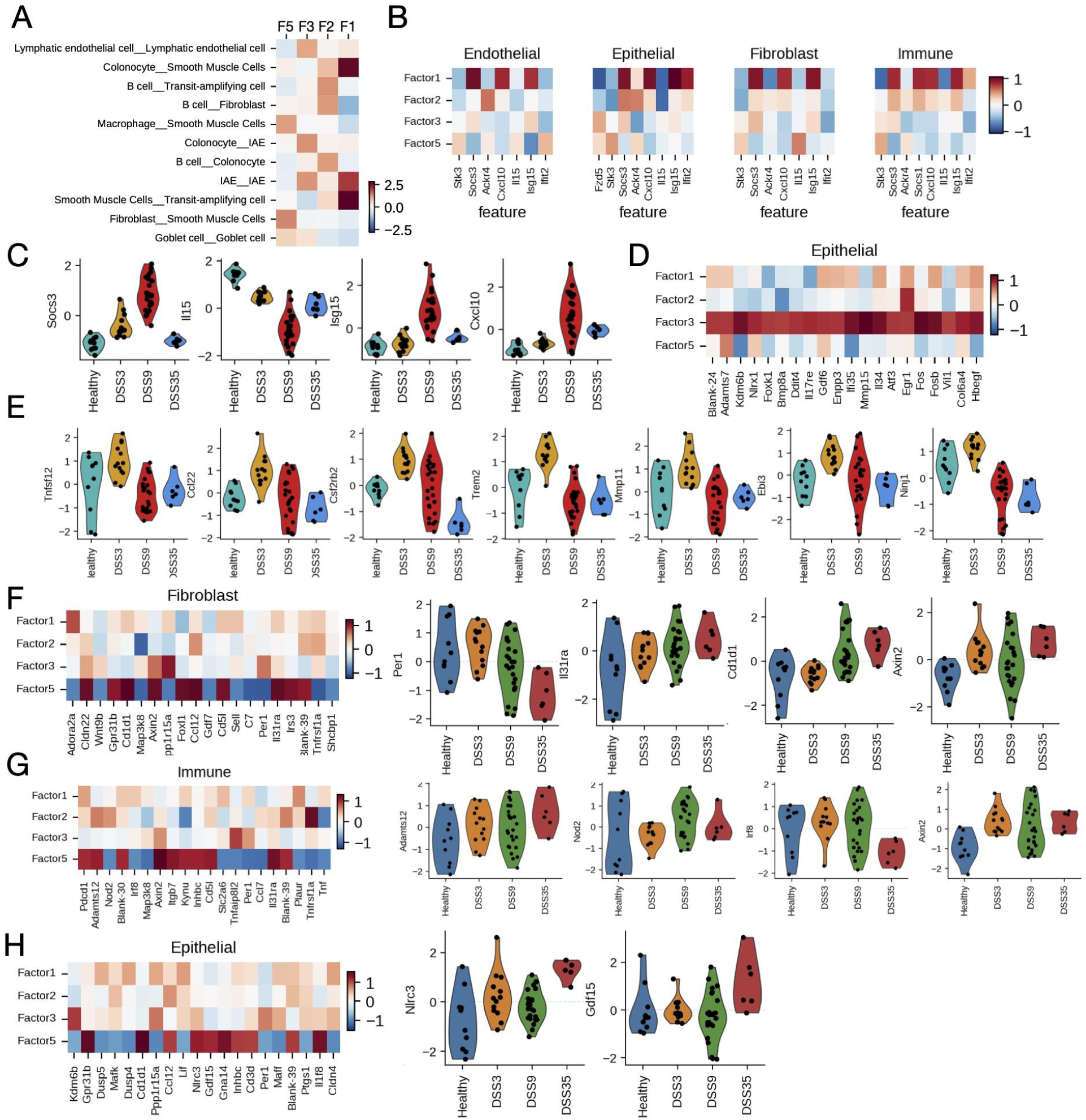
Characterization of spatiotemporal programs of gut damage and recovery. **A.** Top three spatial weights of each factor (summarized as F). **B.** Gene weights of the top two JAK-STAT responsive genes across endothelial, epithelial, fibroblasts and immune cell-types and factors. **C.** Distribution of log-normalized pseudobulked gene expression in epithelial cells of genes belonging to the progressive or peak inflammatory class. **D.** Gene weights across factors of the top 20 genes in Factor 3 and epithelial cells. **E.** Distribution of log-normalized pseudobulked gene expression in immune cells of genes belonging to the early damage response. **F-H** Top 20 genes associated with persistent damage in fibroblasts (F), immune (G) and epithelial cells (H). Right panels show the pseudobulked gene expression of some of the top genes.

